# AR-V7 exhibits non-canonical mechanisms of nuclear import and chromatin engagement in Castrate-Resistant Prostate Cancer

**DOI:** 10.1101/2021.06.03.446940

**Authors:** Seaho Kim, Mohd Azrin Bin Jamalruddin, CheukMan Cherie Au, Eiman Mukhtar, Luigi Portella, Adeline Berger, Daniel Worroll, David S. Rickman, David M. Nanus, Paraskevi Giannakakou

**Author notes:** co-first authors with equal contribution (Seaho Kim, Mohd Azrin Bin Jamalruddin, CheukMan Cherie Au). Address correspondence: Paraskevi Giannakakou, Weill Cornell Medical College, 1300 York Avenue C610C, Department of Medicine, Division of Hematology and Medical Oncology, New York, NY 10021.

## Abstract

Expression of the AR splice variant, AR-V7, in prostate cancer is correlated with poor patient survival and resistance to AR targeted therapies and taxanes. Currently, there is no specific inhibitor of AR-V7, while the molecular mechanisms regulating its biological function are not well elucidated. Here we report that AR-V7 has unique biological features that functionally differentiate it from canonical AR-fl or from the second most prevalent variant, AR-v567. First, AR-V7 exhibits fast nuclear import kinetics via a pathway distinct from the nuclear localization signal dependent importin-α /β pathway used by AR-fl and AR-v567. We also show that the dimerization box domain, known to mediate AR dimerization and transactivation, is required for AR-V7 nuclear import but not for AR-fl. Once in the nucleus, AR-V7 is transcriptionally active, yet exhibits unusually high intranuclear mobility and transient chromatin interactions, unlike the stable chromatin association of liganded AR-fl. The high intranuclear mobility of AR-V7 together with its high transcriptional output, suggest a Hit-and-Run mode of transcription. Our findings reveal unique mechanisms regulating AR-V7 activity, offering the opportunity to develop selective therapeutic interventions.

## Introduction

Androgen receptor (AR) remains a critical therapeutic target in the treatment of metastatic castration resistant prostate cancer (CRPC), due to overactive AR signaling (Feldman and Feldman, 2001). Next generation AR inhibitors targeting either androgen biosynthesis (abiraterone acetate) or AR ligand binding (enzalutamide) have shown improved clinical outcomes including survival. However, these new therapies are not curative due to the development of resistance (Watson et al., 2015). Expression of active AR splice variants (AR-Vs) which re-activate AR transcriptional program in CRPC (Antonarakis et al., 2016; Maughan and Antonarakis, 2015) is one of key drivers in disease progression and is believed to be one mechanism of resistance to abiraterone and enzalutamide. Structurally, the majority of AR-Vs lack the ligand-binding domain (LBD), which is the target of most AR-targeted therapies, and are constitutively active in the nucleus driving AR-signaling (Uo et al., 2018; Watson et al., 2015).

Among more than twenty alternatively spliced AR variants identified to date, AR-V7, which arises from cryptic exon inclusion, is the most prevalent variant in CRPC followed by the exon-skipping AR-v567. Expression of AR-V7 has been clinically associated with adverse patient outcomes including increase rates of metastases and inferior survival rates, and resistance to current standard of care treatment with abiraterone, enzalutamide and taxane chemotherapy (Antonarakis et al., 2014; Antonarakis et al., 2017; Guo et al., 2009; Hornberg et al., 2011; Hu et al., 2009; Maughan and Antonarakis, 2015, Robinson, 2015 #80; Rizzo et al., 2021; Robinson et al., 2015; Tagawa et al., 2018). Together these data suggest that AR-V7 is a driver of CRPC progression and a desirable therapeutic target. However, the exact mechanism(s) underlying AR-V7 oncogenic functions are not well understood. Chromatin immunoprecipitation, transcriptomic and epigenetic studies have identified AR-V7 cistromes and target genes, both distinct and shared with AR-fl, as well as splicing factors that drive AR-V7 production, in an effort to elucidate potentially unique to AR-V7 regulatory mechanisms (Cao et al., 2014; Hu et al., 2009; Li et al., 2013; Melnyk et al., 2020; Xu et al., 2015).

Previously we showed that AR-fl binds microtubules (MTs) via the hinge domain and uses them as tracks for fast nuclear import (Thadani-Mulero et al., 2014; Zhu et al., 2010). Taxanes stabilize MTs and inhibit AR signaling by impairing AR-fl nuclear import and subsequent activation of target genes (Antonarakis et al., 2017; Darshan et al., 2011; Thadani-Mulero et al., 2014; Zhu et al., 2010). Similar to AR-fl, the hinge-containing AR-v567 binds MTs and is sensitive to taxane treatment. In contrast, the hinge-less AR-V7 does not bind MTs, conferring taxane resistance in xenografts and patients with CRPC (Thadani-Mulero et al., 2014, Tagawa et al., 2018).

In this study, we set out to investigate the mechanisms mediating AR-V7 nuclear import and its subnuclear biophysical properties in association with chromatin to identify unique, targetable biological features.

## Results

### AR-V7 exhibits fast nuclear import kinetics in a microtubule- and importin α /β-independent pathway

Nuclear translocation is a pre-requisite for the transcriptional activity of AR-fl and all other nuclear hormone receptors. Nuclear import is mediated by a conserved bipartite nuclear localization signal (NLS) motif, which in AR-fl and AR-v567 is comprised of parts of exons 3 and 4. AR-V7 lacks exon 4, which is the second half of the canonical bipartite NLS, and although its cryptic exon 3 has been implicated in NLS reconstitution (Chan et al., 2012; Guo et al., 2009; Hu et al., 2009), the canonical NLS motif of AR-V7 is compromised. Yet, AR-V7 is constitutively localized to the nucleus in both cell lines and clinical samples, (Guo et al., 2009; Hu et al., 2009; Sun et al., 2010; Watson et al., 2010), indicating efficient nuclear import. To measure basal nuclear import kinetics of the AR-Vs (AR-V7, AR-v567 and AR-fl), plasmids encoding each GFP-tagged AR were microinjected into the nuclei of AR-null PC3 cells and nuclear translocation kinetics was monitored by live-cell time-lapse confocal microscopy (Figure 1A-1B). For each protein we calculated the extent and rate of nuclear import by quantifying the % nuclear GFP-AR protein in single cells over time (Figure 1C). AR-fl remained largely in the cytoplasm under basal condition, exhibiting ∼20% nuclear accumulation which remained steady over the duration of the experiment, indicating very low basal nuclear import kinetics. AR-v567 reached a maximum of ∼50% by 90 min. In contrast, AR-V7 exhibited fast nuclear import kinetics, reaching ∼50% nuclear accumulation within the first 25 min and ∼75% nuclear accumulation by 90 min (Figure 1B-1C and S1A). These data suggested that AR-V7 exhibited the highest basal nuclear import rates, likely via a more efficient nuclear import mechanism. It is established that canonical AR-fl utilizes the classical importin α /β nuclear import mechanism where the importin-α binds to the NLS of AR protein followed by importin β binding, forming a trimeric (cargo-NLS/ importin-α / importin-β) complex in the cytoplasm which enters the nucleus through the nuclear pore complex (NPC) using the Ran-GTP (Pemberton and Paschal, 2005). To identify the mechanisms mediating AR-V7 nuclear import, we examined the involvement of the MT-transport system and the importin-α /β -Ran-GTP pathway (Darshan et al., 2011; Jenster et al., 1993; Kaku et al., 2008; Thadani-Mulero et al., 2014; Zhou et al., 1994; Zhu et al., 2010). We analyzed the translocation kinetics of each variant by live-cell time-lapse imaging using chemical probes that disrupt MTs (docetaxel) or importin-β (importazole) (Figure 1D and S1B-S1D). In agreement with our published data, AR-fl readily translocated to the nucleus upon addition of the synthetic AR ligand R1881, while perturbation of MTs with docetaxel (DTX), abrogated this effect (Thadani-Mulero et al., 2014). Importazole (IPZ), inhibited the R1881-induced AR-fl nuclear translocation, confirming the role of importin-β in the canonical AR-fl nuclear import pathway (Kaku et al., 2008). Similarly, both DTX and IPZ inhibited AR-v567 nuclear import identifying that AR-v567 shares the same pathway of nuclear import with AR-fl (Figure 1D and S1C). In contrast, neither DTX nor IPZ had an effect on AR-V7 nuclear localization, indicating that nuclear import of AR-V7 is both MT and importin-β independent (Figure 1D and S1D).

**Figure 1.**
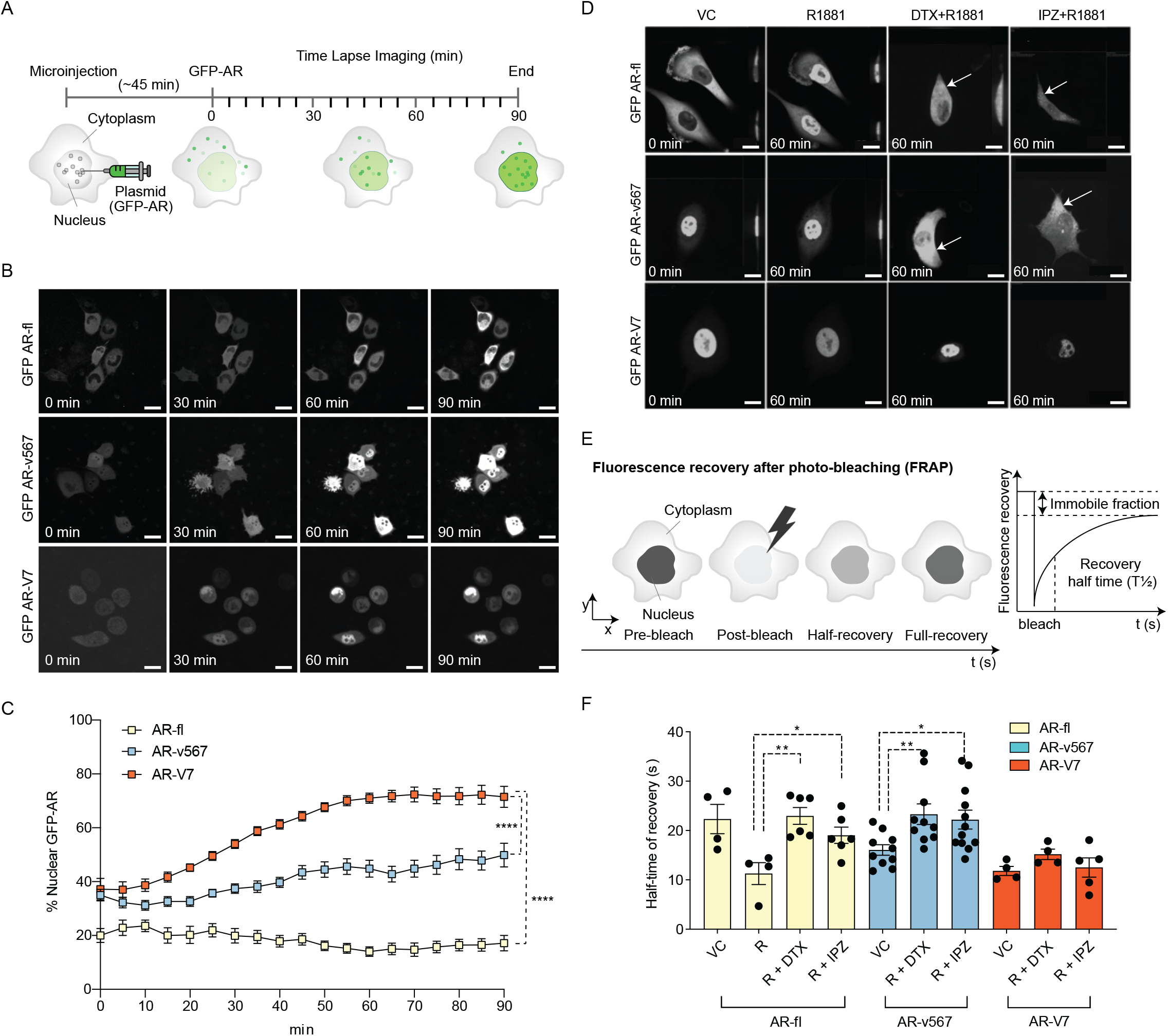
AR-V7 exhibits fast nuclear import kinetics independently of microtubules or the importin-*α /β* pathway, unlike AR-fl or AR-v567. **A**. Experimental design. Plasmids encoding GFP-tagged AR-fl, AR-v567, or AR-V7 were microinjected into the nuclei of the AR-null PC3 cells. As soon as GFP-tagged proteins were detected in the cytoplasm (∼45 min post micro-injection), nuclear translocation kinetics were monitored by live-cell time-lapse confocal microscopy at 5 min intervals for a total of 90 min. **B**. Representative time lapse images showing subcellular localization of each GFP-tagged AR protein. Scale bar, 10 m. **C**. Quantitation of % nuclear GFP-AR protein in single cells (n=3-10 cells/condition/timepoint). **D**. Effect of MT and importin-inhibitors on AR-fl, AR-v567 and AR-V7 nuclear localization. M12 prostate cancer cells stably expressing GFP-tagged AR-fl or AR-v567 or AR-V7 were treated as indicated and subjected to live-cell time lapse imaging. R1881: synthetic androgen; DTX: docetaxel, MT-stabilizing drug; IPZ: importazole, importin-β inhibitor. Representative images are shown. Arrows point to cells with cytoplasmic GFP-AR-fl or GFP-AR-v567. Scale bar, 10µm. **E**. Schematic overview of Fluorescence Recovery After Photobleaching (FRAP) assay and its quantitative output. **F**. Effect of MT and importin-β inhibitors on AR-fl, AR-v567 and AR-V7 nuclear translocation kinetics following FRAP. T1/2 times in sec are shown for each respective protein (n=4-12 cells/ condition). Data represent Mean ± SEM, p-value (*p<0.05, **p<0.01, (****p<0.0001) was obtained using unpaired two-tailed t-test.

To quantify nuclear translocation kinetics of AR proteins in response to treatment, we performed fluorescence recovery after photobleaching (FRAP) analysis in the M12 human metastatic prostate cancer cells stably expressing each GFP-tagged AR protein (Thadani-Mulero et al., 2014). The half-time of recovery (T½; defined as the time required for fluorescence intensity to reach 50% of its pre-bleach intensity) were then calculated and used as a read-out of nuclear import dynamics (Figure 1E). Treatment with R1881 accelerated AR-fl nuclear import by decreasing the T½ from 23 to 11 sec while addition of DTX or IPZ significantly attenuated T½ to 23 and 19 sec, respectively (Figure 1F and S1E). The nuclear recovery of AR-v567, which retains the MT-binding hinge region, was also significantly impaired by DTX or IPZ treatment (T½ 23 and 22 sec respectively). In contrast, AR-V7 nuclear import kinetics were much faster than those of unliganded AR-fl (T½11 vs 23 sec) and were not affected by DTX or IPZ. To determine the involvement of the actin cytoskeleton in AR-V7 nuclear import, we treated PC3 cells microinjected with GFP-tagged AR plasmids with the actin-depolymerizing agent cytochalasin D (Cyto D) and identified that there was no effect on the nuclear import of AR-V7 nor in that of AR-fl or AR-V567 (Figure S1F).

### AR-V7 nuclear import requires active transport via the nuclear pore complex and is partially dependent on Ran-GTP activity

Most nuclear import pathways involve the small GTPase Ran, which catalyzes the release of cargo protein from importin in the nucleus. As AR-V7 does not use importin-α /β pathway for nuclear import, we set out to determine whether it requires active transport via interaction with the nucleoporins, nuclear pore complex components that mediate transport of proteins larger than 40 kD. Thus, we incubated cells with wheat germ agglutinin (WGA), a well-established inhibitor of nucleoporin-mediated nuclear transport (Finlay et al., 1987; Whitehurst et al., 2002; Yoneda et al., 1987) and identified that WGA resulted in cytoplasmic sequestration of GFP-AR-V7 (Figure S2A), suggesting that interaction with nucleoporins is required for AR-V7 nuclear import. Next, to investigate whether AR-V7 depends on Ran-GTP for nuclear import, we quantified the nuclear fraction of GFP-tagged AR-fl, AR-v567, or AR-V7 proteins in the presence of the catalytic Ran-GTP mutant (mCherry-fused Ran Q69L; GTP hydrolysis deficient mutant). Our data identified that AR-fl and AR-v567 nuclear import was inhibited in the presence of the catalytic Ran-GTP mutant (Figure 2A); while, AR-V7 nuclear import was partially inhibited, as evidenced by AR-V7 localization in both nucleus and cytoplasm (Figure 2A; solid arrows). Quantitation of the percent nuclear signal of each AR protein identified significant decrease in nuclear localization for all proteins in the presence of the catalytic mutant RanQ69L (Figure 2B). Similar results were observed when HEK293T cells were transiently co-transfected with GFP-AR-fl, GFP-AR-v567 or GFP-AR-V7 and mCherry-tagged Ran Q69L (Figure S2B-S2C).

**Figure 2.**
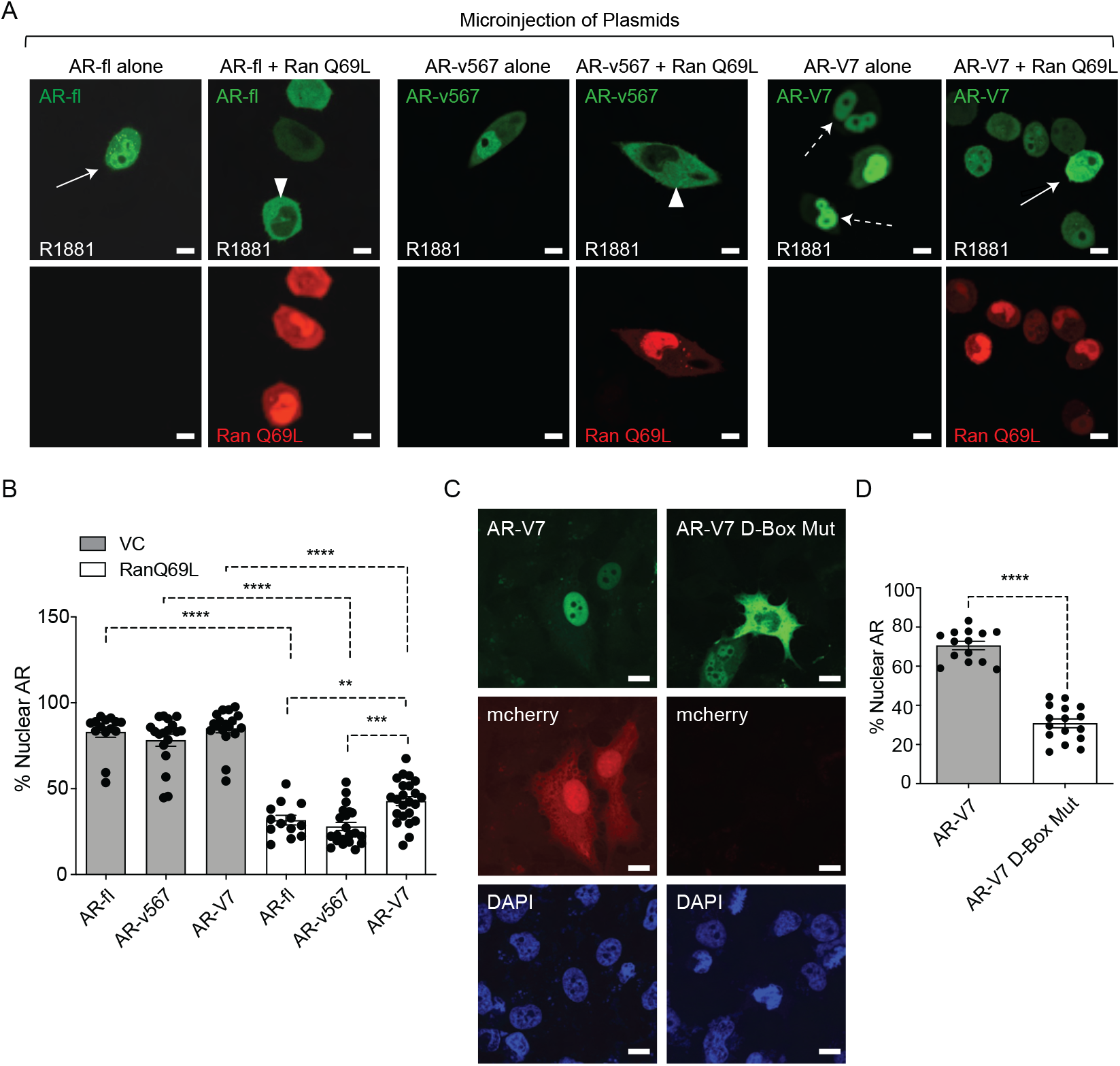
AR-V7 nuclear import requires Ran-GTP-mediated transport via the nuclear pore complex and is impaired upon mutation of the dimerization box domain (D-Box). **A**. Ran-GTP is required for AR-V7 nuclear import. Plasmids encoding GFP-tagged AR-fl, AR-v567, or AR-V7 were co-microinjected with plasmid encoding the catalytic mutant Ran-GTP (mCherry-tagged RanQ69L), into the nuclei of the AR-null PC3 cells. Cells expressing both tagged proteins were subjected to live-cell time lapse imaging. Cells expressing AR-fl were treated with R1881 (10 nM) to induce nuclear import. Representative confocal microscopy images show nuclear localization of the mCherry-tagged RanQ69L and varying subcellular localization of each GFP-tagged AR protein. Solid arrow: cell with both cytoplasmic and nuclear AR proteins; arrowheads: cells with cytoplasmic AR proteins; dashed arrow: cell with nuclear AR proteins. Scale bar, 10 µm. **B**. Graphic display of % Nuclear AR across n>10 per condition. **C-D**. AR-V7 nuclear import is impaired upon mutation of the dimerization box domain (D-box). PC3 cells stably expressing ARE-mCherry reporter were transfected with GFP-AR-V7 or GFP-AR-V7 D-box mutant (A596T, S597T). **(C)** Representative images and **(D)** quantitative results are shown (n>10 cells per condition). Data represent Mean ± SEM, p-value (**p<0.01, ***p< 0.001, ****p<0.0001) was obtained using unpaired two-tailed t-test.

### AR-V7 nuclear import is impaired upon mutation of the dimerization box domain (D-Box)

Androgen-regulated gene expression requires AR-fl receptor dimerization in the nucleus mediated by the zinc finger (D-box) domain, prior to DNA binding. AR-V7 transcriptional activity has been shown to depend on the D-box domain shown to mediate AR-V7 homodimerization in the nucleus (Centenera et al., 2008; Xu et al., 2015). To determine the potential impact of D-box domain on AR-V7 nuclear import which is a pre-requisite for its transcriptional activity, we generated a construct encoding GFP-AR-V7 containing two established, functionally-inactivating D-box mutations (A596T, S597T) and transiently transfected PC3 cells stably expressing the ARE-mCherry reporter in order to measure ARE activity in single cells (Azeem et al., 2017). Cells transfected with GFP-AR-V7 alone had an average of 71% nuclear AR-V7 with concomitant mCherry expression indicating ARE-transactivation. In contrast, cells transfected with GFP-AR-V7-D-box mutant not only did they become transcriptionally inactive, but they also exhibited extensive cytoplasmic enrichment with significantly lower nuclear localization as compared to GFP-AR-V7 (from 71% in AR-V7 to 31% in AR-V7-D-box mutant; p<0.0001) (Figure 2C-2D). Interestingly, the same D-box mutations had no effect on AR-fl nuclear import (Figure S2D) while, as expected, they abrogated its transcriptional activity (not shown). Together, these results indicate that AR-V7 nuclear import is impaired upon mutation of D-box, identifying a novel, variant-specific function for this conserved domain.

### AR-V7 drives ligand-independent fractional nuclear translocation of AR-fl with no evidence of heterodimerization

In CRPC, AR-fl is often co-expressed with AR-V7 in patient tumors and it has been suggested that active AR signaling in castrate conditions is partially due to AR-V7 heterodimerization with AR-fl resulting in its nuclear translocation in the absence of ligand (Cao et al., 2014; Xu et al., 2015). Such a mechanism implies a common pathway of nuclear import for the AR-V7/AR-fl heterodimer. To explore this possibility, we microinjected mCherry-AR-fl and GFP-AR-V7 plasmids either individually or together into PC3 cells and quantified their respective nuclear accumulation (Figure 3A-3B). Under basal conditions we observed low AR-fl nuclear localization which was increased by 5-fold in the presence of ligand (12% *vs* 66%, p<0.0001); while AR-V7 alone was predominantly localized in the nucleus (79%). When both AR-V7 and AR-fl were co-expressed in the same single cell, in the absence of ligand, AR-fl nuclear localization was increased by 2-fold compared to AR-fl expressed alone (24% vs 12%, respectively, p<0.001). Similar results were observed following transient co-transfection (Figure S3A-S3B). To determine whether AR-V7 might form a heterodimer with unliganded AR-fl in the cytoplasm driving the latter into the nucleus, we used the importin-β inhibitor importazole (IPZ), which as we showed earlier inhibits AR-fl nuclear import (Figure 1 and S1B-S1E). Our results identified that when AR-V7 and AR-fl were co-expressed in the same cell, IPZ inhibited the nuclear localization of AR-fl only; while it had no effect on AR-V7 (Figure 3C), consistent with the results obtained with IPZ treatment when each protein was expressed alone. These results suggest lack of physical interaction between AR-fl and AR-V7 in the cytoplasm in agreement with recently published proteomics data (Chen et al., 2018) and lack of co-precipitation between AR-fl and AR-V7 (Guo et al., 2009).

**Figure 3.**
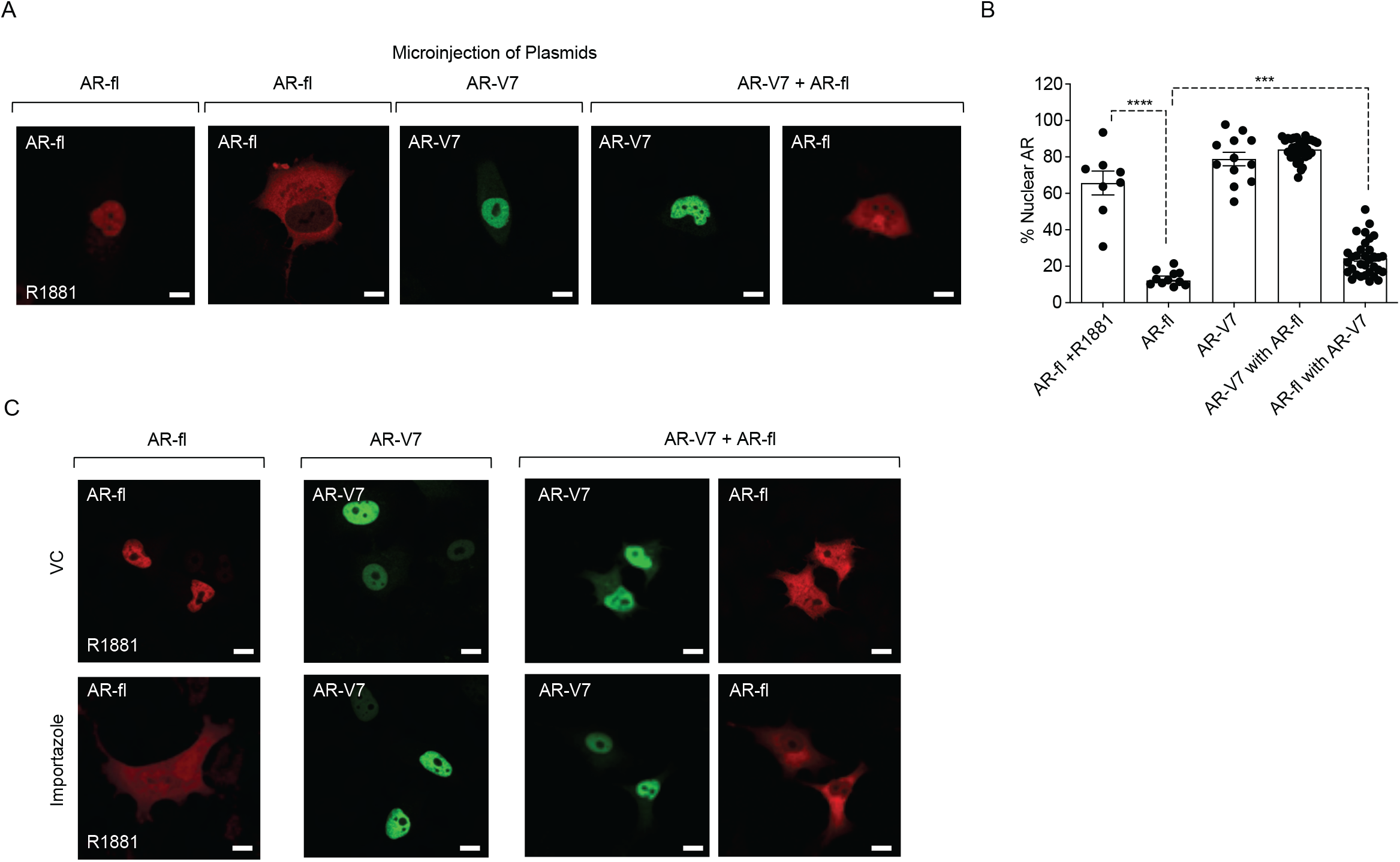
AR-V7 drives nuclear translocation of AR-fl in the absence of ligand. **A-B**. Plasmids encoding mCherry-AR-fl or GFP-AR-V7 were micro-injected in PC3 cells. **A-B**. Representative microscopic images (scale bar, 10 µm) and % nuclear AR is shown. Data represent Mean ± SEM with n>10 cells per condition, p-value (***p< 0.001,****p<0.0001) was obtained using unpaired two-tailed t-test. **C**. PC3 cells were transfected with mCherry-AR-fl, or GFP-AR-V7, and cells were treated 10 nM R1881 or 50 µM Importazole, as indicated. Representative confocal microscopy images are shown for each condition. Scale bar,10 µm.

To investigate whether AR-V7 transcriptional output mediates the fractional AR-fl nuclear translocation, we introduced the DNA-binding domain (DBD) mutation (A573D) known to inactive AR-fl, into AR-V7 (AR-V7-A573D). We found that the A573D mutation abrogated AR-V7 transcriptional activity compared to unaltered AR-V7 (Figure 4A-4C), as evidenced by the decrease in ARE-mCherry reporter activity and the decreased expression of the endogenous AR-V7 target gene, FKBP5. When the transcriptionally inactive AR-V7-A537D mutant was co-expressed with AR-fl, we observed a similar increase in nuclear AR-fl in the presence of mutant or unaltered AR-V7 as compared to baseline levels (31% vs 39 vs 19%, respectively, p<0.0001) (Figure 4D-4E).

**Figure 4.**
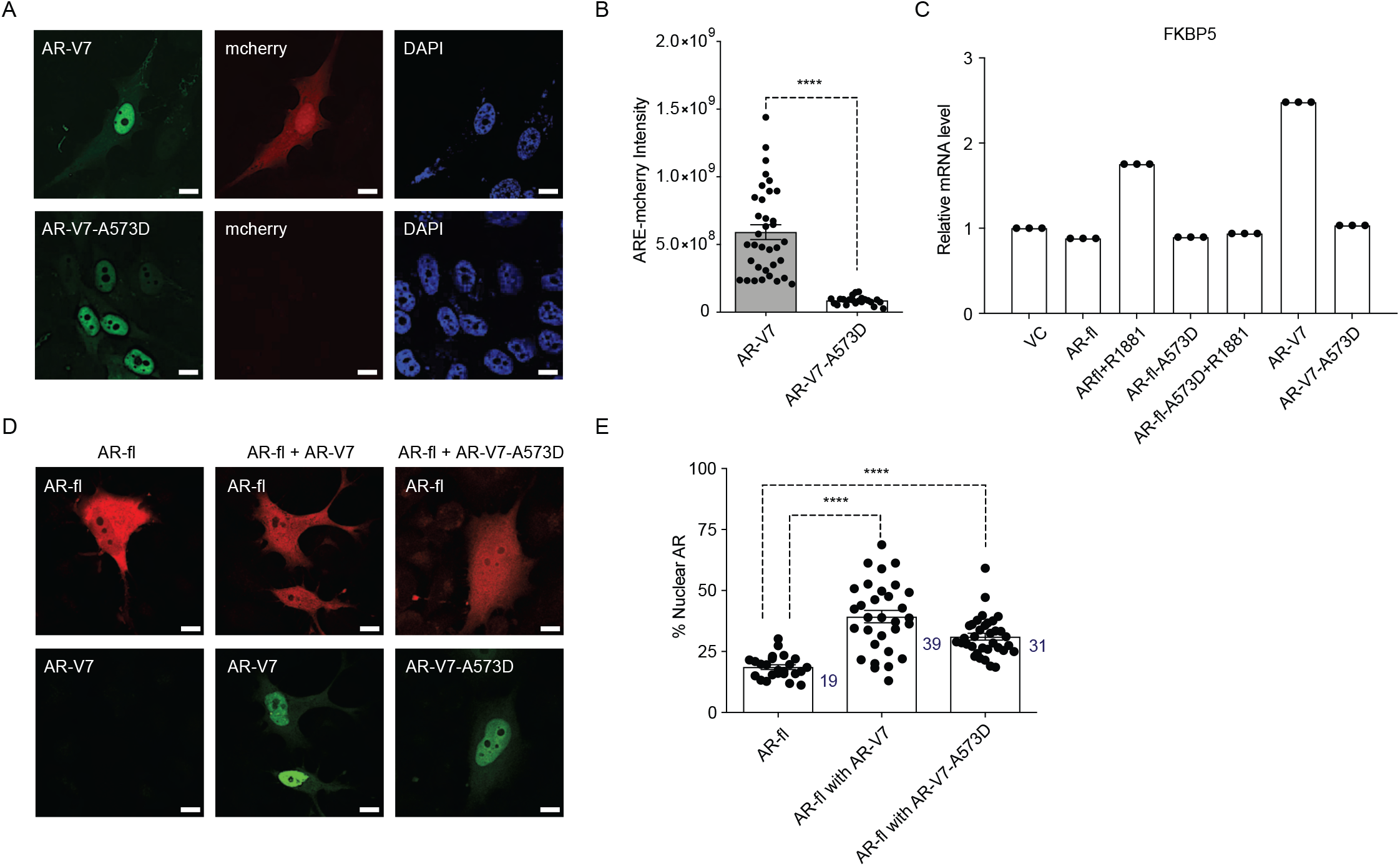
DBD mutation abrogates AR-V7 transcriptional activity. **A-B**. PC3 cells stably expressing ARE-mCherry reporter were transfected with the indicated plasmids for 48 hrs and the expression of GFP protein with concomitant mCherry protein was analyzed by confocal imaging. Representative images and quantitative results are shown (n>10 cells per condition). Scale bar, 10µm. **C**. RT-qPCR for the endogenous FKBP5 mRNA was quantified in PC3 cells after transfection of indicated plasmids. Data with AR-fl A573D are included as a control (n=3). **D**. PC3 cells were transfected with mCherry-AR-fl or GFP-AR-V7 or GFP-AR-V7-A573D and imaged by confocal microscopy (Scale bar, 10µm) and **E**. % nuclear AR protein was quantified (n>23 cells per condition). Data represent Mean ± SEM, p-value (****p<0.0001) was obtained using unpaired two-tailed t-test.

### AR-V7 exhibits high subnuclear mobility kinetics and short chromatin residence time

It is well established that agonist-bound AR-fl is transcriptionally active and relatively immobile in the nucleus; while, antagonist-bound AR-fl is highly mobile and transcriptionally inactive (Farla et al., 2004; Farla et al., 2005). AR-V7, on the other hand, is transcriptionally active in the absence of agonist-binding, however, the exact mechanism underlying its nuclear activity is not known. Thus, we investigated the exchange rate of AR-V7 with chromatin using FRAP and confocal live cell imaging in cells transfected with GFP-tagged AR-V7 or AR-fl as a control. FRAP analysis of AR-fl identified slow the fluorescence recovery of ligand-bound nuclear AR was significantly delayed compared to unliganded AR (T½ ∼8s vs 3s, respectively, p<0.0001) (Figure 5A-5C), suggesting prolonged chromatin residence time of ligand-bound AR, in agreement with reports on AR and other nuclear hormone receptors including the glucocorticoid, estrogen and progesterone receptors (Farla et al., 2004; Farla et al., 2005; Klokk et al., 2007). In contrast, the fluorescence recovery of nuclear GFP-AR-V7 was very fast compared to R1881-bound AR-fl (T½ ∼4s and 8s, respectively, p<0.0001) (Figure 5A-5C), indicating that AR-V7 exhibits short chromatin residence time, despite being transcriptionally active.

**Figure 5.**
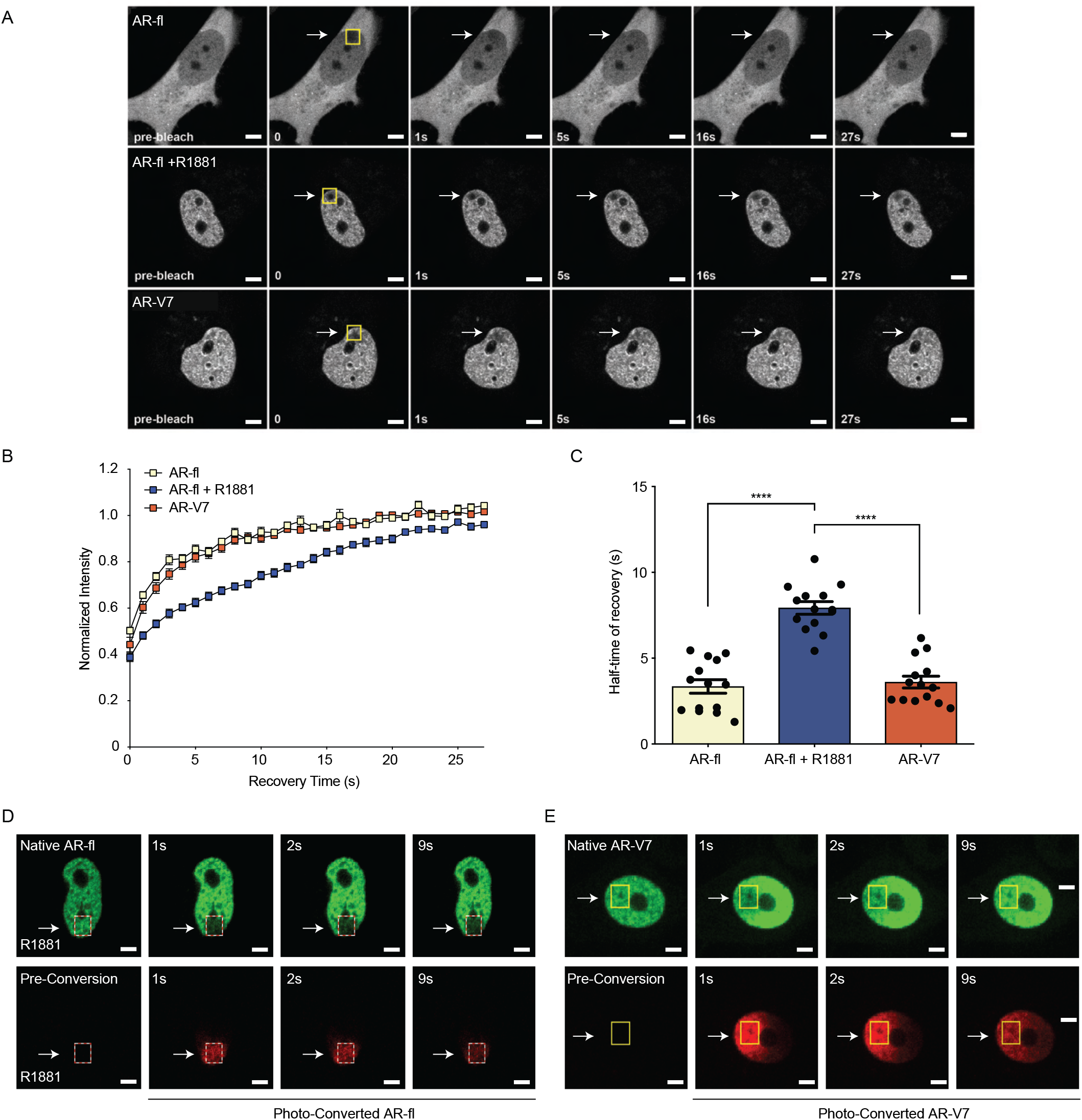
AR-V7 exhibits high intranuclear mobility compared with liganded AR-fl. **A**. FRAP was performed in PC3 cells transiently expressing GFP-AR-fl (in the absence or presence of 10 nM R1881) or GFP-AR-V7. FRAP was monitored at 1s intervals. Representative images of cells at select time points are shown. **B**. Kinetics of proteins recovery after photobleaching at 1s intervals are graphically displayed, n=14. **C**. Graphic display of half-time of recovery (T 1/2) in seconds (s) for each condition, n=14. **D-E**. FRAP was performed in PC3 cells transiently expressing photoconvertible mEos4b-AR-fl or mEos4b-AR-V7 protein. Cells were imaged at 1s intervals to monitor fluorescence recovery of the non-converted proteins (Green) and nuclear distribution of photo-converted proteins (Red). Scale bar: 10µm. Data represent Mean ± SEM, p-value (****p<0.0001) was obtained using unpaired two-tailed t-test.

To better understand the kinetics of subnuclear mobility of each protein we generated photo-convertible mEos4b-AR-fl or mEos4b-AR-V7 and followed them by live cell imaging (Paez-Segala et al., 2015). Photo-conversion of AR-fl in a small sub-nuclear region, led to a change in the fluorophore from green (un-converted) to red (converted), which was monitored by time-lapse imaging at 1s intervals (Figure 5D, dotted box). Our data identified that AR-fl remained within the confines of the photoconverted area, without fluorescence recovery by unconverted AR (green) suggesting stable and prolonged chromatin binding (Figure 5D). On the other hand, photo-converted AR-V7, in < 1s started moving outside the photoconverted area (Figure 5E, yellow box), repopulating the entire nucleus in less than 9s. This high intranuclear mobility of AR-V7 was accompanied by rapid fluorescence recovery of un-converted-AR-V7 (green) (Figure 5E). These data reveal for the first time a sharp distinction between the chromatin exchange rates and intranuclear mobility of AR-V7 and canonical ligand-bound AR-fl, despite their comparable transcriptional output. To investigate any potential nuclear interactions of both proteins when co-expressed in the same cell, we co-microinjected mCherry-AR-fl and GFP-AR-V7 in the nuclei of PC3 cells and monitored their respective recovery after photobleaching. We observed a similar pattern of intranuclear kinetics for each protein, when co-expressed in the same sub-nuclear area, as the kinetics observed when each protein was expressed alone with fast AR-V7 fluorescence recovered vs. slow recovery of ligand-bound AR-fl (Figure S4A-S4C).

### DNA binding mutation abrogates AR-V7 transactivation and accelerates nuclear mobility kinetics

To resolve the conundrum between the high nuclear mobility of AR-V7 and its high transcriptional activity, we introduced the A573D DBD mutation into AR-V7 or AR-fl expression plasmids and introduced them into PC3 cells. We detected a significant increase in the mobility of the mutant AR-fl-A573D compared to AR-fl, in the presence of ligand, with T ½ of 4s *vs* 17s, respectively, p<0.0001 (Figure 6A and S5). No difference was observed in the absence of ligand, suggesting that immobilization of the ligand-bound AR-fl is mediated by the DNA-binding domain of AR-fl. Surprisingly, we found that the already high intranuclear mobility of AR-V7 was further accelerated by the DBD mutation with recovery T ½ for AR-V7-A573D at 2s vs. 4s for AR-V7, p<0.0001 (Figure 6B). These data suggested that DNA-binding mediates the transient chromatin interactions exhibited by AR-V7. Next, we examined the effect of the DBD mutation on the transcriptional output of AR-fl and AR-V7, using ARE-mCherry expression as a transcriptional readout in single cells, or target gene mRNA expression by RT-qPCR in cell populations. Our results revealed that the A573D mutation abrogated transcriptional activity in both, ligand-bound AR-fl and AR-V7, as evidenced by the significant decrease in ARE-mCherry expression (Figure 6C-6D). Together, these data couple DNA binding with nuclear mobility kinetics and AR-V7 transactivation.

**Figure 6.**
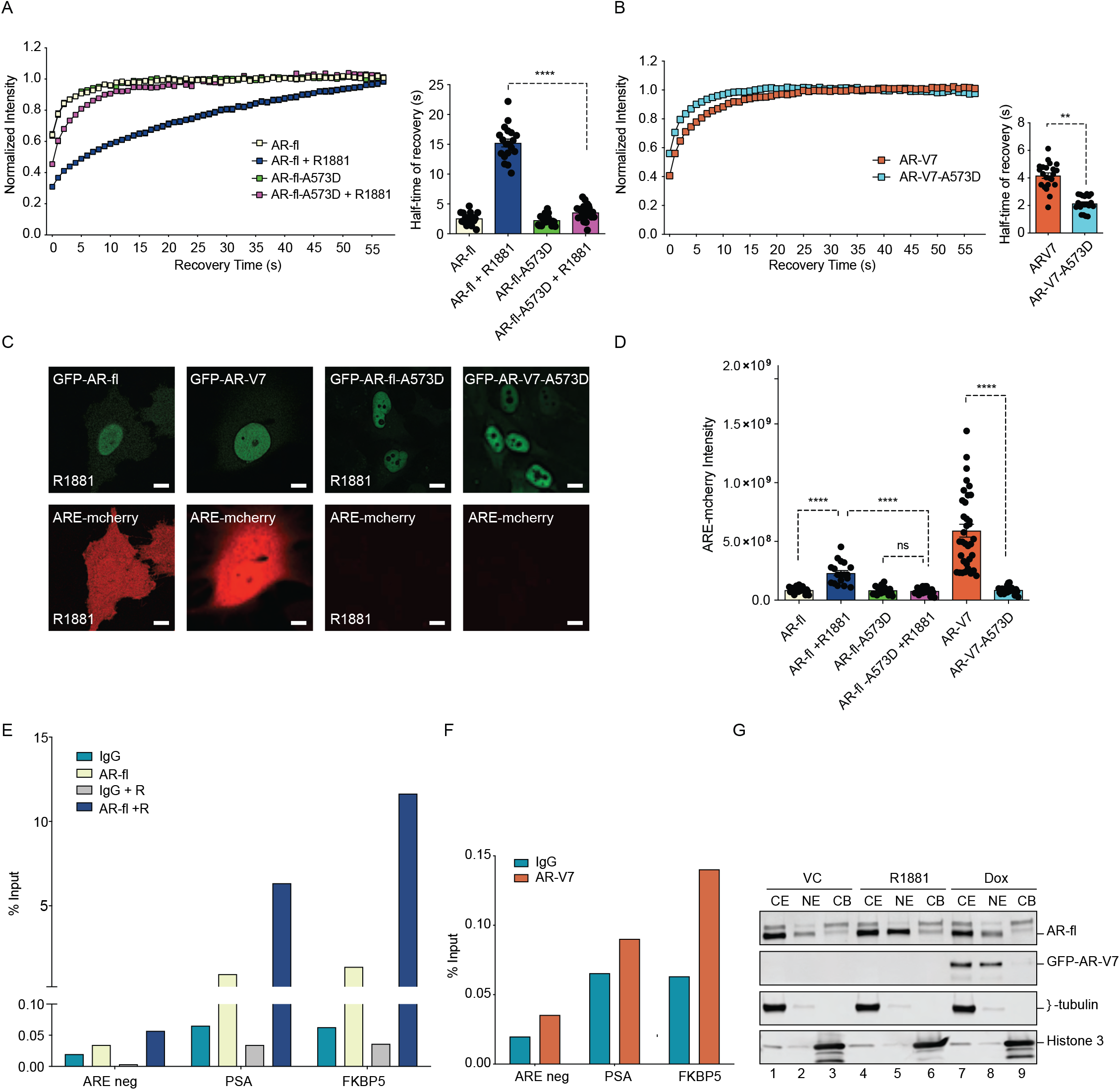
DNA-binding mutation increases the intranuclear mobility and abrogates the transcriptional activity of AR-fl and AR-V7. **A-B**. FRAP was performed in PC3 cells transiently expressing GFP-AR-fl or GFP-AR-V7 or their respective DBD mutants (A573D) Kinetics of protein recovery after photobleaching are graphically displayed and their half-time of recovery, is obtained for each condition, n>14. **C**. PC3 cells stably expressing ARE-mCherry reporter were transfected with indicated plasmids, in the presence or absence of ligand (10 nM R1881). Representative images of each condition are shown. **D**. quantitation of mCherry fluorescence intensity in single cells (n>17). **E-F**. The binding of **(E)** AR-fl or **(F)** AR-V7 on the enhancer of PSA or FBKP5 in 22RV1 was analyzed by ChIP-QPCR assay. Cells in charcoal stripped media were treated with vehicle or 10 nM R1881 for 24hrs. **G**. Immunoblot for AR-fl and AR-V7 following subcellular fractionation CE, cytosolic extract; NE, nuclear extract, CB, chromatin-bound nuclear extract. Histone H3 and β -tubulin were used as controls for the fractionation. Data represent Mean ± SEM, p-value (**p<0.01, ****p<0.0001) was obtained using unpaired two-tailed t-test.

To investigate whether the high intranuclear mobility of AR-V7 is due to its reduced occupancy rate on target AREs on chromatin, we performed chromatin immunoprecipitation (ChIP) for AR-V7 or AR-fl in 22Rv1 cells with endogenous expression of both proteins. ChIP-qPCR data showed that R1881 increased the occupancy of AR-fl on the enhancer regions of PSA and FKBP5 (Figure 6E). In contrast, AR-V7 showed very low occupancy rate on PSA and FKBP5 enhancer (∼0.01 % input) compared to ligand-bound AR-fl occupancy (6∼12 % Input). (Figure 6F). To corroborate these data we performed sub-cellular fractionation of C4.2:GFP-AR-V7 cells expressing endogenous AR-fl and doxycycline-inducible AR-V7 (Figure 6G). As expected, R1881 enhanced both the nuclear AR-fl (NE) and chromatin-bound AR-fl (CB) fractions (Figure 6G). In contrast, minimal if any AR-V7 was detected in the chromatin-bound fraction, consistent with its low chromatin occupancy rate and high subnuclear mobility kinetics.

## Discussion

Inhibition of androgen receptor signaling remains the cornerstone of contemporary therapeutic strategies for patients with metastatic CRPC. Reactivation of AR signaling is a hallmark of CRPC, largely mediated by the nuclear activity of the AR splice variant AR-V7 (Antonarakis et al., 2014; Antonarakis et al., 2017; Guo et al., 2009; Hornberg et al., 2011; Hu et al., 2009; Maughan and Antonarakis, 2015).

In this study, we identify that AR-V7 nuclear import does not use the canonical NLS-dependent importin-α /β pathway, in contrast to AR-fl and AR-v567. Earlier findings suggested that the C-terminal cryptic exon 3 (CE3) domain of AR-V7 might mediate its nuclear import due to it similarity with the second bipartite NLS of AR-fl. (Chan et al., 2012). The same study, also showed that a synthetic truncated AR-V7 lacking the CE3 domain, did not bind importin-α, yet was transcriptionally active in the nucleus. These data not only show the CE3 motif is dispensable for nuclear import, but they are also consistent with our results where pharmacologic inhibition of importin-β, has no effect on AR-V7 nuclear localization confirming an NLS-independent mechanism of AR-V7 nuclear entry. The molecular weight of AR-V7 protein together with its cytoplasmic sequestration upon nucleoporin inhibition suggest that AR-V7 requires active transport via the NPC, likely using an alternative nuclear transporter (Figure S2A). The β -like importin family molecules, known to mediate nuclear import for proteins without a classical bipartite NLS by recognizing an alternative nuclear import signal, are potential candidates for AR-V7 nuclear import pathway (Chen et al., 2017).

Surprisingly, we found that the same inactivating mutations introduced in the D-box domain (A596T and S597T), identified in patients with androgen insensitivity syndrome (Centenera et al., 2008), had a different effect on AR-fl compared to AR-V7. It is well established that the D-box domain is important for AR dimerization in the nucleus, which occurs prior to DNA binding, and that mutations in this domain impair nuclear AR dimerization and activation of target genes (van Royen et al., 2012). Using bimolecular fluorescence complementation (BiFC) assay, Xu et al. showed that these D-box mutations inhibited AR-V7 homo- and hetero-(with AR-fl) dimerization in the nucleus (Xu et al., 2015). In contrast, our results (Figure 2C) revealed that D-box mutant AR-V7 is transcriptionally inactive because it is sequestered in the cytoplasm, suggesting that the D-box domain may be important for the nuclear translocation of AR-V7, which is distinct from its function in AR-fl.

Analysis of single PC3 cells co-expressing tagged AR-FL and AR-V7, showed significant increase in AR-fl nuclear localization, in the absence of ligand (Figure 3C, S3) in agreement with recent reports (Cao et al., 2014). Together these results suggested a potential physical interaction between the two proteins in the cytoplasm and a shared mechanism of nuclear import. To test this hypothesis, we used IPZ to inhibit importin-β mediated AR-fl nuclear import and observed no effect on AR-V7, implying that AR-V7 and AR-fl use independent nuclear import pathways and that likely there is no physical interaction between the two proteins (Figure 3C). Along these lines, previous studies showed that AR-V7 does not co-precipitate with AR-fl and that they do not form a physical complex in the nucleus (Chen et al., 2018; Guo et al., 2009). Interestingly, when the transcriptionally inactive AR-V7-A537D mutant was co-expressed with AR-fl, we observed a similar fractional increase in AR-fl nuclear translocation, in the absence of ligand (Figure 4D, E) implicating non transcriptional mechanisms. It should be noted that the mere localization of AR-fl in the nucleus in castrate conditions does not imply AR signaling activation. Taken together our own findings and published results, we posit that there is no evidence that AR-V7 mediated fractional increase in nuclear AR-fl contributes to the overall activity of AR-V7 in driving castrate-independent growth in PC.

This study also revealed unique nuclear biology of AR-V7, distinct from that of ligand-bound AR-fl, suggesting a distinct mode of transcriptional action. The chromatin binding dynamics of nuclear hormone receptors (glucocorticoid, progesterone, estrogen, androgen) has been closely correlated with their respective transcriptional output (Farla et al., 2004; Farla et al., 2005; Fletcher et al., 2000; Klokk et al., 2007; McNally et al., 2000; Stenoien et al., 2001). It is well established that AR-V7 is constitutively active in the nucleus having largely overlapping cistromes and target genes with canonical AR-fl (Cato et al., 2019). However, its mode of transcription is not well elucidated. Herein, using live cell imaging and photo-conversion to quantify subnuclear dynamics of AR proteins, we show that ligand-bound AR-fl is transcriptionally active and exhibits low intranuclear mobility, prolonged residence time on chromatin and high occupancy rates on promoter AREs (Figure 5 and 6). Conversely, AR-V7 is transcriptionally active, yet exhibits unusually high intranuclear mobility and transient chromatin interactions with low occupancy rates. The high intranuclear mobility of AR-V7 together with its high transcriptional output, suggest a Hit-and-Run model of transcription, where a transcription factor (TF) transiently binds a DNA sequence to regulate target genes (the “hit”), and before vacating the site (the “run”) recruits secondary TFs which form stable complex at the regulatory site that sustain a stable long-term effect (Charoensawan et al., 2015). Interestingly, the Hit-and-Run is often applied to transcriptional repressors, where gene silencing does not necessarily require continuous TF residence on chromatin (Shah et al., 2019). Importantly, a recent study showed that AR-V7 functions as a transcriptional repressor in CRPC, preferentially binding several co-repressors compared to AR-fl (Cato et al., 2019). Finally, the reported diversity of AR-V7 regulated transcriptomes across patients with CRPC, which likely results from the cell-context specific AR-V7 cistromes (Chen et al., 2018) is compatible with a Hit-and-Run mode of transcription that allows for fast adaptation to environmental cues.

## Acknowledgements

We thank Dr. Alison North (the Bio-Imaging Resource Center at Rockefeller University) for assistance with the photo-conversion study. We are grateful to Dr. Karl-Henning Kalland and Dr. Waqas Azeem from University of Bergen, Norway (ARE-mCherry reporter), Dr. David Root from Broad Institute (pCW57.1 tet-on vector, Addgene Plasmid # 41393), Dr. Jay Brenman from University of North Carolina at Chapel Hill (pmCherry-C1-RanQ69L, Addgene plasmid # 30309) for provision of reagents and valuable experimental advice. This work was supported by grants from the US NIH T32 CA203702 (S. Kim), US NIH T32 CA062948 (C.C. Au), R01CA137020 (P. Giannakakou), R21CA216800 (P. Giannakakou), R01CA228512 (P. Giannakakou); R01CA179100 (P. Giannakakou and D.S. Rickman); and from the Department of Defense W81XWH-17-1-0162 (A. Berger).

## Author contributions

S. K, M.A.B.J, C.C.A with equal contribution of Conceptualization, Data curation, Formal analysis, Funding acquisition, Validation, Investigation, Methodology, Writing - original draft, Writing - review and editing; E.M, L.P, A.B, D.W Investigation, Methodology; D.S.R Resources, Methodology; D.M.N, Resources, Supervision; P.G, Conceptualization, Resources, Formal analysis, Supervision, Funding acquisition, Methodology, Writing-original draft, Project administration, Writing - review and editing

## Materials and Methods

### Cell lines

PC3, LNCaP, C4-2, HEK-293T, and 22Rv1 cell lines were obtained from the ATCC. We generated he C4-2 cell line (tet-on GFP-AR-V7), which stably expresses tetracycline-inducible GFP-AR-V7 by infecting the lentiviral construct (detailed information in construct section). The stable M12 cell lines expressing GFP-tagged AR-fl, AR-v567, or AR-V7 were described previously (Thadani-Mulero et al., 2014)

### Antibodies and reagents

Primary antibodies were: rabbit anti-AR-N-terminal (AR-N-21), rabbit monoclonal anti-AR-V7 (31-1109-00, RevMab), rabbit monoclonal anti-AR-C-terminal (ab52615, abcam), rabbit anti-actin (A2066, SIGMA), rabbit anti-beta tubulin (ab6046), rabbit anti-Histone H3 (ab1791). Docetaxel (Toxotere), importazole, cytochalasin D were obtained from Sigma. Wheat Germ Agglutinin (WGA), Alexa Flour 594 Conjugate (W11262) was purchased from molecular probes.

### Plasmid constructions

The following plasmids pmCherry-AR-fl, pEGFP-C1-AR-fl, pEGFP-C1-AR-v567, and pEGFP-AR-V7 (Thadani-Mulero et al., 2014) were used for transfection or microinjection into the cell nuclei. The DNA-binding domain (DBD) mutant at A573D of AR-fl and AR-V7 in the pEGFP-C1 backbone were generated by site-directed mutagenesis using the Q5 site-Directed Mutagenesis Kit (New England BioLabs). The dimerization box (D-box) mutant at A596T/S597T of AR-V7 in the pEGFP-C1 backbone was also generated by the same mutagenesis approach.

The photoconvertible mEos4b-C1 backbone (Addgene plasmid # 54812) was a gift from Dr. Michael Davidson (Paez-Segala et al., 2015). We subcloned AR-fl, AR-v567, or AR-V7 in mEos4b-C1 to generate N-terminally tagged-photoconvertible AR constructs (mEos4b-AR-fl, mEos4b-AR-v567, and mEos4b-AR-V7) used for live cell imaging. Doxycycline inducible GFP-tagged AR-V7 was generated by sub-cloning of GFP-AR-V7 into the lentiviral pCW57.1 tet-on vector (a gift from Dr. David Root, Addgene Plasmid # 41393) using Gateway cloning (Invitrogen). The construct containing mCherry-tagged GTP-hydrolysis defective Ran mutant, pmCherry-C1-RanQ69L (Addgene plasmid # 30309), was a gift from Dr. Jay Brenman (Kazgan et al., 2010) and used for live cell imaging. ARE-reporter vector CS-GS241B-mCHER-LV152 with mCherry fluorescent reporter signal was a gift from Dr. Karl-Henning Kalland (Azeem et al., 2017). Using this vector, we generated lentiviral particles to infect PC3 cells. Stable PC3 cells harboring CS-GS241B-mCHER-LV152 were generated by hygromycin (500 µg/ml) selection and used for ARE-mCherry reporter assay.

### Live-cell imaging, FRAP, and photo-conversion analysis

Live cell imagining was carried out on cells either microinjected or transfected with the plasmids described above. Cells were grown on No. 1.5 coverglass mounted on 35 mm MatTek dish and cultured in RPMI 1640 supplemented with 5% charcoal-stripped FBS, 25 mM HEPES, 2 mM sodium pyruvate, and 2 mL L-glutamine. Live cell imaging and Fluorescence recovery after photobleaching (FRAP) were carried out on a Zeiss LSM 700 confocal microscope equipped with an on-stage live-cell chamber (Tokai Hit, Shizuoka, Japan). Photo bleaching in the region of interest was carried out with the 405-nm laser at maximum power for 3 iterations. A single z-section was imaged before and at time intervals. The normalized intensity of region of interests and the half-time of recovery required for the fluorescence intensity to reach 50% of its pre-bleach intensity (T1/2) were obtained using Zeiss Zen software. Photo-conversion imaging analysis was performed in PC3 cells plated on 35 mm MatTek dish after transient transfection with photoconvertible AR constructs descried above. PC3 cells transfected with mEos4b-AR-fl was further incubated with 10 nM R1881 for 1h before the photo-conversion. Briefly, region of interest at nuclei of cells was photo-converted by applying 405 nm laser (60% power for 400 ms) using the Mosaic system (Andor, Oxford Instruments, UK) equipped with spinning disk confocal microscope (Zeiss/Perkin-Elmer) at Bio-imaging resource center at the Rockefeller University. The time-lapse images were captured in two different channels (for green 491 nm laser, 525-50 nm filter; for red 561 nm laser, 620-60 nm filter) before and after photo-conversion and images acquisition was performed with MetaMorph software.

### ARE-mCherry reporter assay

PC3 cells stably expressing the ARE-reporter vector with mCherry fluorescent reporter signal was plated on 35 mm MatTek dish. GFP-tagged AR-fl or AR-V7 was microinjected in the nuclei of cells. The synthetic androgen R1881 was added to the cells microinjected with GFP-AR-fl construct. After overnight incubation, GFP-AR and ARE-reporter mCherry signal were imaged using Zeiss scanning confocal microscope.

### Quantitative real-time PCR

For relative quantitation of AR target genes, quantitative real-time PCR was performed on 100 ng input RNA using Power SYBR green RNA-to-Ct 1 step kit (Applied Biosystems) and primers specific for PSA (F: 5’-ACGCTGGACAGGGGGCAAAAG, R: GGGCAGGGCACATGGTTCACT), FKBP5 (F: 5’-GCGGAGAGTGACGGAGTC, R: 5’-TGGGGCTTTCTTCATTGTTC), and ACTIN (F: 5’-CCTCCCTGG AGAAGAGCTA, R: 5’-CCAGACAGCACTGTATTGG). Relative quantitation was used to determine fold change in expression levels by the comparative Ct method.

### Chromatin Immunoprecipitation

22RV1 cells were trypsinized and crosslinked in 1% formaldehyde media for 10 minutes at room temperature and quenched for 8 min using 125 mM glycine. Nuclear extracts were collected and sonicated for 10 minutes to obtain 300bp chromatin fragments (Diagenode Bioruptor Pico). Equal volumes of sheared chromatin were immunoprecipitated with rabbit AR-V7 antibody (RevMab 31-1109-00), rabbit AR antibody (abcam 52615), or rabbit IgG control (Santa Cruz). After extensive washing, crosslinking was reversed, and DNA fragments were purified using Macherey-Nagel kit (740609). Q-PCR amplification was performed using the ABI 7500 fast system (Fast SYBR Green 4385612 Applied Biosystems) and the relative standard curve method in a 96-well format. Primers used: FKBP5e: F-GGT TCC TGG GCA GGA GTA AG; R-AAC GTG GAT CCC ACA CTC TC; PSAe: F-TGG GAC AAC TTG CAA ACC TG; R-GAT CCA GGC TTG CTT ACT GT; AREneg: F-GCT GAT TCA ATT ACC TCC CAG AA; R-AGT TTG GGA CAG ACG GGA AA. The input chromatin for each sample was analyzed at 4 concentrations (serial dilutions) to generate a standard curve per primer pair and per 96-well plate. The sheared chromatin was diluted 1/6 before being used for Q-PCR. All reactions were run in triplicate.

### Subcellular fractionation and Western Blot analysis

C4.2 cell line expressing tet-inducible GFP-AR-V7 was treated with either 10 nM R1881 for AR nuclear translocation or 1 ug/ml doxycycline for GFP-AR-V7 induction. Sub-cellular fractionation of cells was performed using the subcellular protein fractionation kit (Thermo Scientific) and each extract was subjected to immunoblotting with indicated antibodies.

### Statistical analyses

The Student two-tailed t-test was used to determine the mean differences between two groups. P < 0.05 is considered significant. Data were presented as mean ± SEM.

## Supplementary information titles and legends

**Figure S1. Related to Figure 1. AR-V7 exhibits fast nuclear import kinetics independently of microtubules, actin or the importin-α /μ pathway. A**. Plasmid encoding GFP-tagged AR-V7 was microinjected into nuclei of PC3 cells and as soon as GFP was detected in the cytoplasm (∼45 min post micro-injection) the kinetics of GFP-AR-V7 nuclear import were monitored by live-cell time-lapse confocal microscopy at 5 min intervals for a total of 180 min. Representative time lapse images are shown at the indicated time points. Solid arrow: cell with both cytoplasmic and nuclear AR-V7 at time 0; Arrowhead: cells with cytoplasmic only AR-V7 at time 0; Dashed Arrow: cell with primarily nuclear AR-V7 first detected at +50 min after the start of imaging. Enhanced AR-V7 nuclear translocation is observed over time for all cells. Notice that there are cells with already extensive nuclear accumulation of AR-V7 at 0 min, suggesting very fast nuclear import kinetics from the time of microinjection (−45 min). Scale bar, 10 μm. **B-D**. Corresponds to Figure 1D with additional time points. Briefly, M12 prostate cancer cells stably expressing GFP-tagged (B) AR-fl, (C) AR-v567 or (D) AR-V7 were treated as indicated and subjected to live-cell time lapse imaging. R1881: synthetic androgen used to stimulate AR-fl nuclear translocation; DTX: docetaxel, MT-stabilizing drug; IPZ: importazole, importin-β inhibitor. Representative images are shown. Scale bar 10 μm. **E**. Table with T1/2 (half-time recovery) values for each variant (related to Fig. 1F). **F**. PC3 cells were treated with 1 μg/ml cytochalasin D (Cyto D) or vehicle control (VC) for 1 hour at 37°C following plasmid micro-injection into the nuclei of PC3 cells. Cells were then treated with 10 nM R1881 for 4 hrs and subjected to point-scanning confocal microscopy. Representative images showing are shown. Scale bar 10 μm.

**Figure S2. Related to Figure 2. AR-V7 nuclear import requires active transport via the nuclear pore complex, is dependent on Ran-GTP activity and is impaired upon mutation of the dimerization box domain (D-Box). A**. Wheat germ agglutinin (WGA) blocks AR-V7 nuclear import: we incubated cells with WGA, an inhibitor of nucleoporin-mediated nuclear transport and monitored GFP-AR-V7 localization by live cell imaging. WGA kept AR-V7 in the cytoplasm in the presence of doxycycline suggesting that AR-V7 nuclear import requires active transport via the NPC. Representative confocal microscopy images were shown. Red: WGA labeling of membranes; Green: GFP-AR-V7. Nuc: nuclear AR-V; Cyto: cytoplasmic AR-V7. Scale bar represent 10 μm. **B**. (related to Fig. 2B) HEK293T cells were transfected with plasmids encoding GFP-tagged AR-fl, AR-v567 and AR-V7 in the presence of the catalytic mutant Ran-GTP (mCherry-tagged RanQ69L). Nuclear accumulation of each AR variant was calculated. Representative confocal microscopy images (63x magnification) for each condition are shown. Solid arrow: cell with both cytoplasmic and nuclear proteins; arrowheads: cells with cytoplasmic protein; dashed arrow: cell with nuclear protein. Scale bar represent 10 μm. **C**. Graphic display of % Nuclear AR across 30 individual cells per condition. Data represent Mean ± SEM with n>10, p-value (**p<0.01,****p<0.0001) was obtained using unpaired two-tailed t-test. **D**. PC3 cells were transfected with GFP-AR-fl or GFP-AR-fl-D-box mutant (A596T, S597T) and they were treated for 4 hr with 10 nM DHT. There was no effect of the D-box mutations on AR-fl nuclear import.

**Figure S3. Related to Figure 3. AR-V7 drives nuclear translocation of AR-fl in the absence of ligand. A-B**. Plasmids encoding mCherry-AR-fl or GFP-AR-V7 were transfected in PC3 cells. Representative microscopic images (scale bar, 10 μm) and % nuclear AR is shown. Data represent Mean ± SEM with n>10 cells per condition, p-value (**p< 0.01,****p<0.0001) was obtained using unpaired two-tailed t-test.

**Figure S4. Related to Figure 5. AR-V7 intranuclear mobility is not affected by co-expression of ligand-bound AR-fl. A**. FRAP was performed in PC3 cells co-microinjected with mCherry-AR-fl and GFP-AR-V7 followed by treatment with 10 nM R1881 for 2hrs. Representative images from the same single cell co-expressing the two AR proteins are shown. **B**. Kinetics of protein recovery after photobleaching for each protein, when co-expressed in the same single cell, are graphically displayed. The fluorescence intensity in the bleached area was measured and depicted as the normalized recovery. **C**. Half-time of recovery (T1/2) required for the fluorescence intensity to reach 50% of its pre-bleach intensity was compared, n=5. Data represent Mean ± SEM, p*-*value (**p< 0.01) is obtained using unpaired two-tailed t-test.

**Figure S5. Related to Figure 6. DBD mutation increases the intranuclear mobility of liganded-AR-fl and AR-V7. A**. FRAP was performed in the nuclei of PC3 cells following transient expression of GFP-AR-fl or GFP-AR-fl A573D (in the presence of 10 nM R1881) or GFP-AR-V7 or GFP-AR-fl A573D. Representative images of cells at the indicated time points are shown. Arrow heads show the photo-bleaching area. Scale bar, 10 μm.

