## Supplementary Figures for "AR-V7 exhibits non-canonical mechanisms of nuclear import and chromatin engagement in Castrate-Resistant Prostate Cancer"

Figure S1

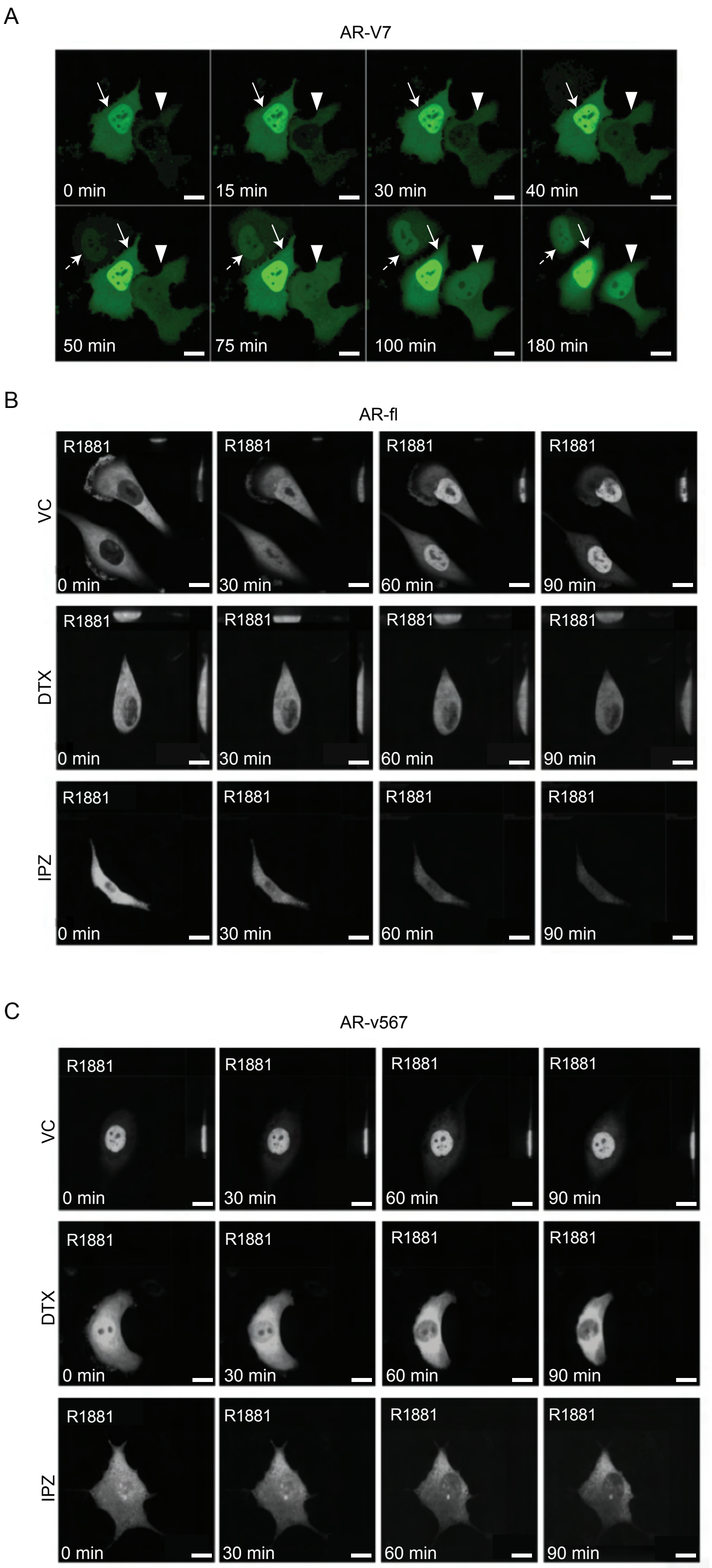


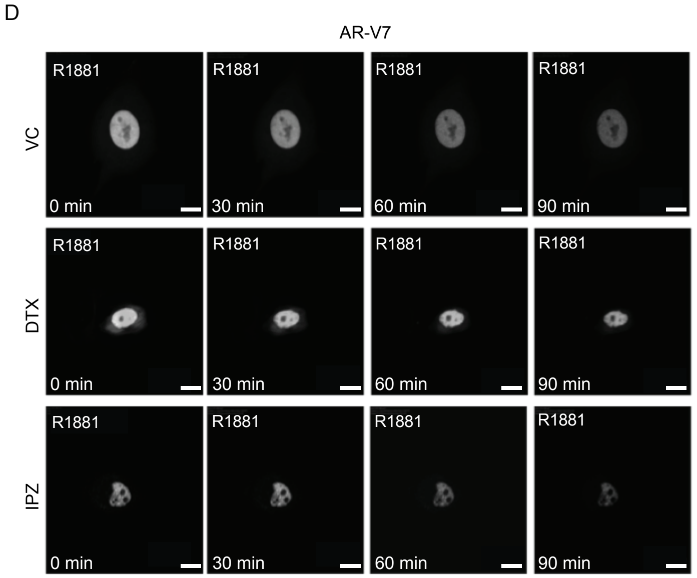


E

**Half-time of recovery (s)**

| | AR-fl | AR-v567 | AR-V7 |
|---|---|---|---|
| VC | 23 | 16 | 11 |
| R | 11 | - | - |
| R + DTX | 23 | 23 | 15 |
| R + IPZ | 19 | 22 | 12 |

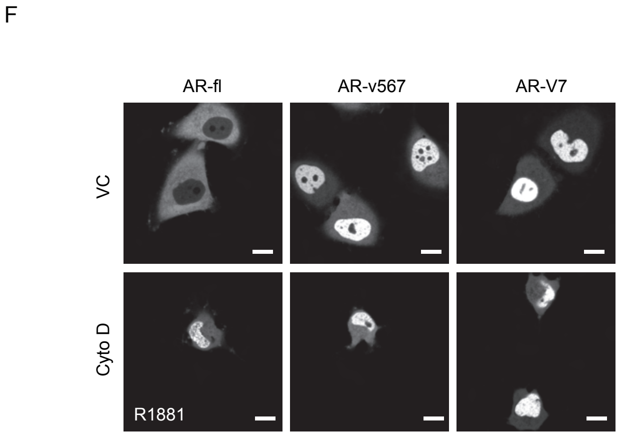


**Figure S1. Related to Figure 1. AR-V7 exhibits fast nuclear import kinetics independently of microtubules, actin or the importin-α/β pathway. A.** Plasmid encoding GFP-tagged AR-V7 was microinjected into nuclei of PC3 cells and as soon as GFP was detected in the cytoplasm (~45 min post micro-injection) the kinetics of GFP-AR-V7 nuclear import were monitored by live-cell time-lapse confocal microscopy at 5 min intervals for a total of 180 min. Representative time lapse images are shown at the indicated time points. Solid arrow: cell with both cytoplasmic and nuclear AR-V7 at time 0; Arrowhead: cells with cytoplasmic only AR-V7 at time 0; Dashed Arrow: cell with primarily nuclear AR-V7 first detected at +50 min after the start of imaging. Enhanced AR-V7 nuclear translocation is observed over time for all cells. Notice that there are cells with already extensive nuclear accumulation of AR-V7 at 0 min, suggesting very fast nuclear import kinetics from the time of microinjection (-45 min). Scale bar, 10 μm. **B-D.** Corresponds to Figure 1D with additional time points. Briefly, M12 prostate cancer cells stably expressing GFP-tagged **(B)** AR-fl, **(C)** AR-v567 or **(D)** AR-V7 were treated as indicated and subjected to live-cell time lapse imaging. R1881: synthetic androgen used to stimulate AR-fl nuclear translocation; DTX: docetaxel, MT-stabilizing drug; IPZ: importazole, importin-  inhibitor. Representative images are shown. Scale bar 10μm. **E.** Table with T1/2 (half-time recovery) values for each variant (related to Fig. 1F). **F.** PC3 cells were treated with 1 μg/ml cytochalasin D (Cyto D) or vehicle control (VC) for 1 hour at 37°C following plasmid micro-injection into the nuclei of PC3 cells. Cells were then treated with 10 nM R1881 for 4 hrs and subjected to point-scanning confocal microscopy. Representative images showing are shown. Scale bar 10 μm.

Figure S2

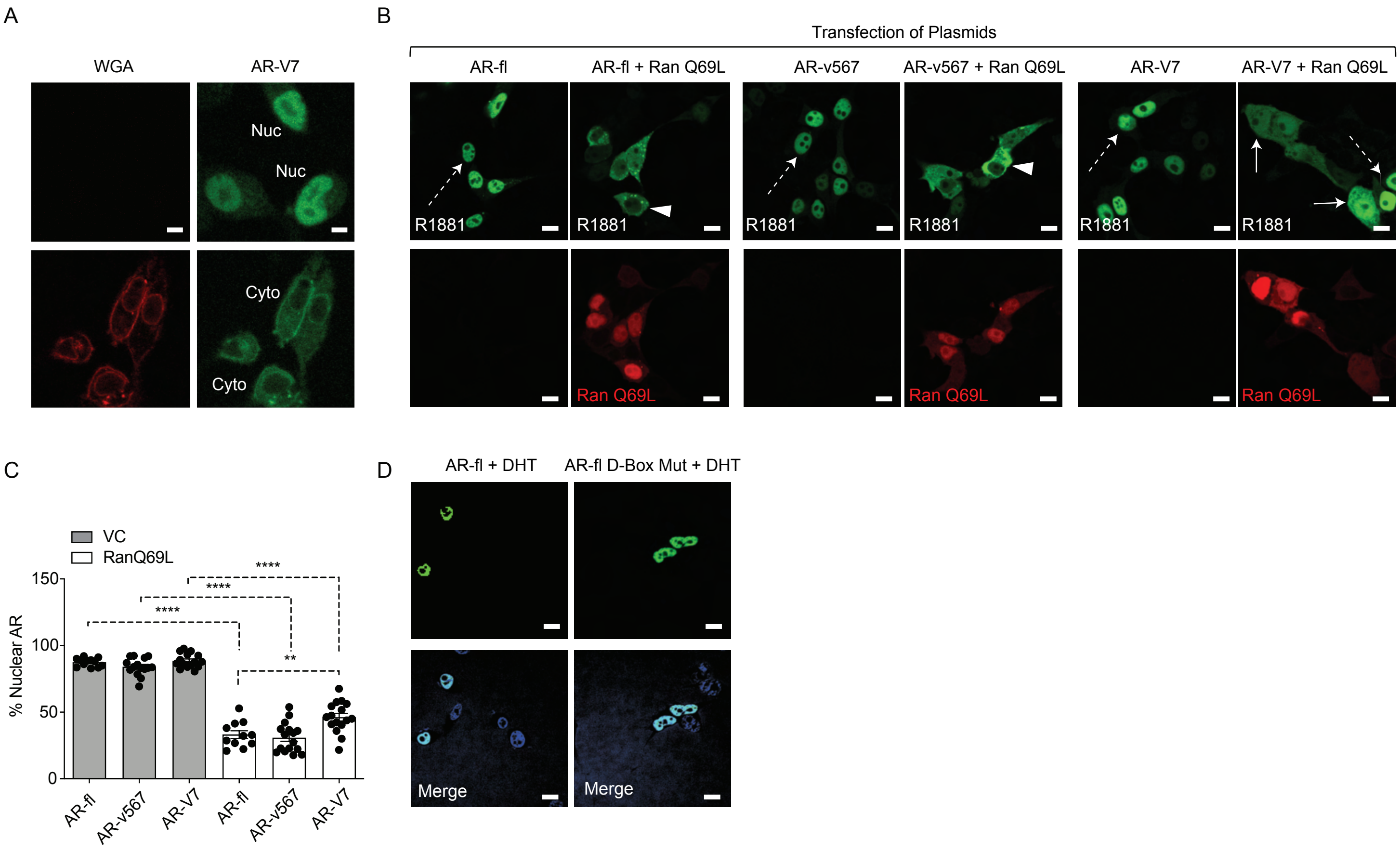


**Figure S2. Related to Figure 2. AR-V7 nuclear import requires active transport via the nuclear pore complex, is dependent on Ran-GTP activity and is impaired upon mutation of the dimerization box domain (D-Box). A.** Wheat germ agglutinin (WGA) blocks AR-V7 nuclear import: we incubated cells with WGA, an inhibitor of nucleoporin-mediated nuclear transport and monitored GFP-AR-V7 localization by live cell imaging. WGA kept AR-V7 in the cytoplasm in the presence of doxycycline suggesting that AR-V7 nuclear import requires active transport via the NPC. Representative confocal microscopy images were shown. Red: WGA labeling of membranes; Green: GFP-AR-V7. Nuc: nuclear AR-V; Cyto: cytoplasmic AR-V7. Scale bar represent 10 μm. **B.** (related to Fig. 2B) HEK293T cells were transfected with plasmids encoding GFP-tagged AR-fl, AR-v567 and AR-V7 in the presence of the catalytic mutant Ran-GTP (mCherry-tagged RanQ69L). Nuclear accumulation of each AR variant was calculated. Representative confocal microscopy images (63x magnification) for each condition are shown. Solid arrow: cell with both cytoplasmic and nuclear proteins; arrowheads: cells with cytoplasmic protein; dashed arrow: cell with nuclear protein. Scale bar represent 10μm. **C.** Graphic display of % Nuclear AR across 30 individual cells per condition. Data represent Mean ± SEM with n>10, p-value (**p<0.01,****p<0.0001) was obtained using unpaired two-tailed t-test. **D.** PC3 cells were transfected with GFP-AR-fl or GFP-AR-fl-D-box mutant (A596T, S597T) and they were treated for 4 hr with 10 nM DHT. There was no effect of the D-box mutations on AR-fl nuclear import.

Figure S3

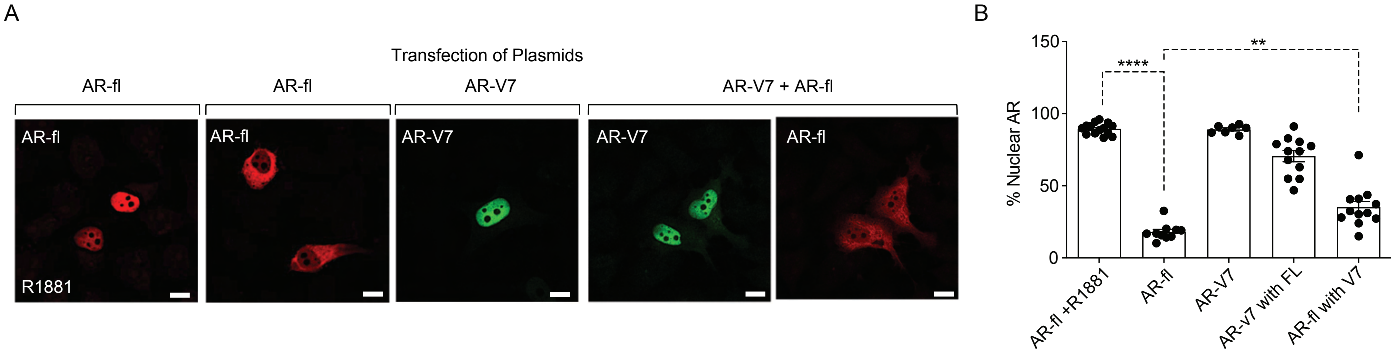


**Figure S3. Related to Figure 3. AR-V7 drives nuclear translocation of AR-fl in the absence of ligand. A-B.** Plasmids encoding mCherry-AR-fl or GFP-AR-V7 were transfected in PC3 cells. Representative microscopic images (scale bar, 10 μm) and % nuclear AR is shown. Data represent Mean ± SEM with n>10 cells per condition, p-value (**p< 0.01,****p<0.0001) was obtained using unpaired two-tailed t-test.

Figure S4

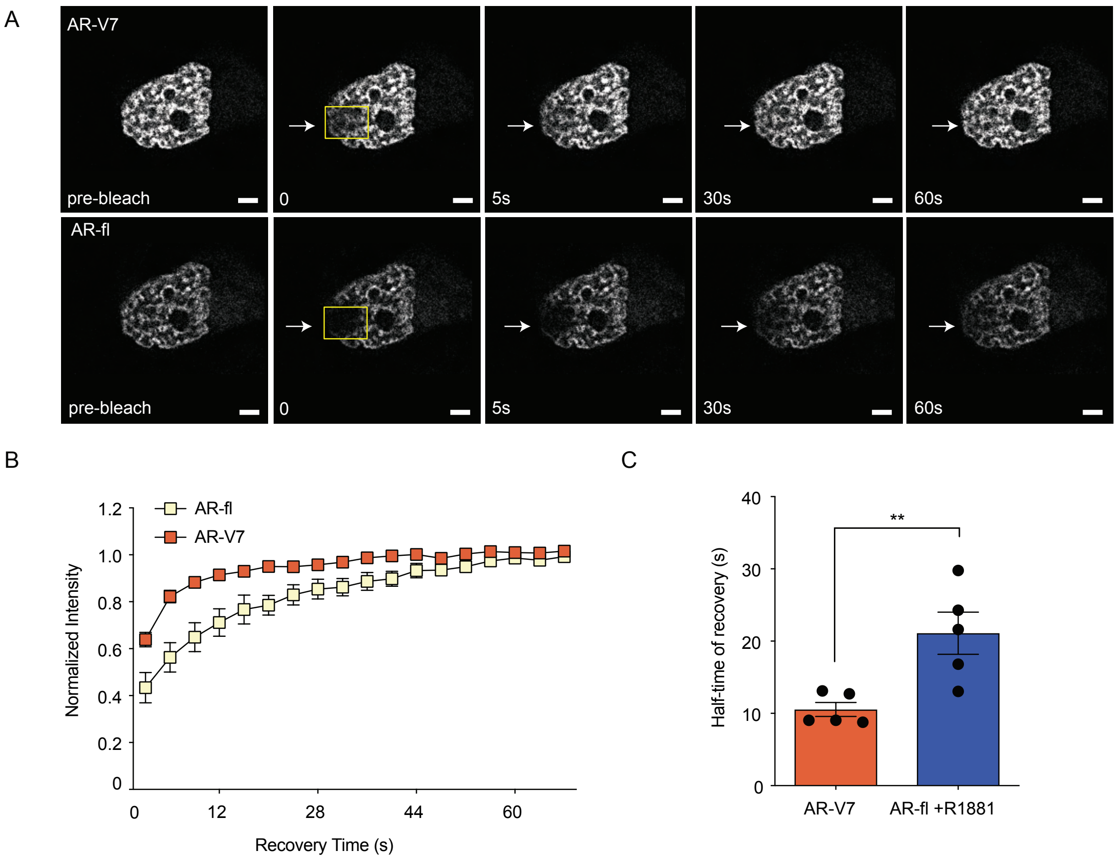


**Figure S4. Related to Figure 5. AR-V7 intranuclear mobility is not affected by co-expression of ligand-bound AR-fl. A.** FRAP was performed in PC3 cells co-microinjected with mCherry-AR-fl and GFP-AR-V7 followed by treatment with 10 nM R1881 for 2hrs. Representative images from the same single cell co-expressing the two AR proteins are shown. **B.** Kinetics of protein recovery after photobleaching for each protein, when co-expressed in the same single cell, are graphically displayed. The fluorescence intensity in the bleached area was measured and depicted as the normalized recovery. **C.** Half-time of recovery (T1/2) required for the fluorescence intensity to reach 50% of its pre-bleach intensity was compared, n=5. Data represent Mean ± SEM, p-value (**p< 0.01) is obtained using unpaired two-tailed t-test.

Figure S5

A

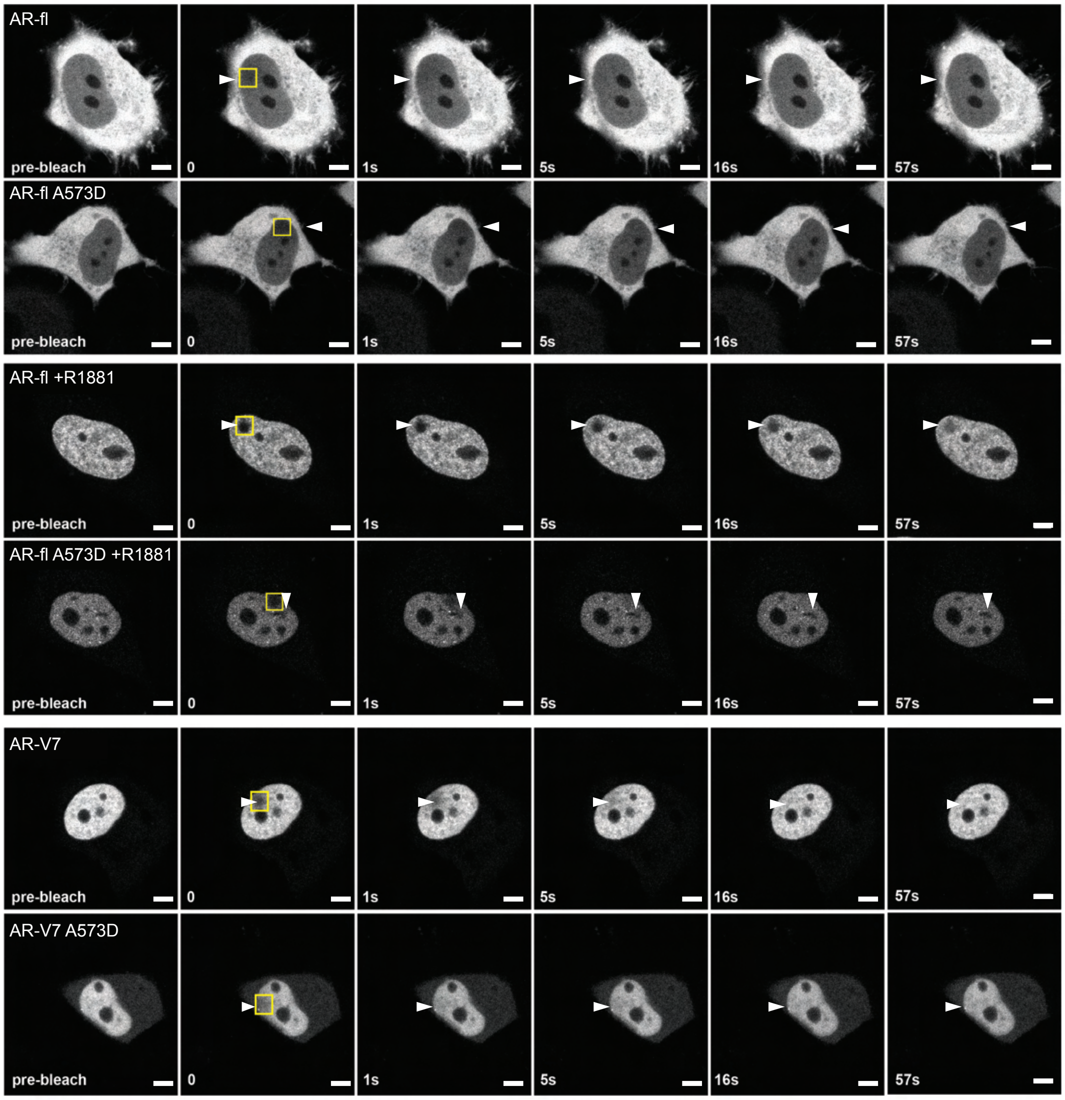


**Figure S5. Related to Figure 6. DBD mutation increases the intranuclear mobility of liganded-AR-fl and AR-V7. A.** FRAP was performed in the nuclei of PC3 cells following transient expression of GFP-AR-fl or GFP-AR-fl A573D (in the presence of 10 nM R1881) or GFP-AR-V7 or GFP-AR-fl A573D. Representative images of cells at the indicated time points are shown. Arrow heads show the photo-bleaching area. Scale bar, 10μm.
